# A new perspective on slow pacemaking in brain and heart

**DOI:** 10.1101/2025.10.30.685563

**Authors:** Arthur Fyon, Oleksandra Pavlova, Nick Schaar, Pietro Mesirca, Amandine Ligot, Marvin Gaillardon, Julien Brandoit, Sofian Ringlet, Alessio Franci, Matteo E. Mangoni, Jochen Roeper, Guillaume Drion, Vincent Seutin, Kevin Jehasse

## Abstract

Slow pacemaking is a key physiological process in specific excitable cells, yet its underlying mechanism remains debated. Here we identify a conserved, voltage-dependent pacemaker current that is essential for slow, regular firing in both midbrain dopaminergic neurons and sinoatrial node myocytes. Conductance-based models incorporating this current reproduce stable pacemaking, requiring a fast-activating, small-amplitude current. This is further confirmed by dynamic-clamp experiments in dopaminergic neurons. Replacing the pacemaker current in a model with a voltage-independent conductance such as the non-selective sodium leak channels fails to sustain slow rhythmicity, highlighting the necessity for an adequate voltage dependence. Our results suggest a novel and shared biophysical mechanism for slow pacemaking in neuronal and cardiac systems.

## Introduction

Pacemaking is a fundamental property of a subset of excitable cells that underpins vital rhythmic functions across biological systems. Pacemaker cells can be classified, based on the firing frequency, as either fast or slow pacemakers. While the mechanism sustaining fast pacemaking is known ^1^, slow pacemaking remains controversial. It has been suggested that it originates from the balanced interplay between several depolarizing and hyperpolarizing currents ^2–6^. The main ionic currents involved in pacemaking are hyperpolarization-activated cyclic nucleotide-gated (HCN encoding *I_H_* in neurons and *I_F_* in cardiac cells) and voltage-gated Ca^2+^ (Ca_v_) and Na^+^ (Na_v_) channels underlying inward currents on the one hand, and A-type (*I_A_*) fast inactivating, ERG-type (*I_Kr_*), or KvLQT (*I_Ks_*) K^+^ channels, underlying outward currents on the other hand. Yet, experimental data shows that blocking or altering any of ion channels driving inward current, do not abrogate slow pacemaking, although both its frequency and regularity can be affected ^3,7–12^. Another hypothesis is that the production of such a precise (coefficient of variation below 5%) and robust low frequency (1-5 Hz on average) pacemaking needs the reliable activation of a pacemaker current (*I_pace_*) during the interspike interval ^1,7,13–16^. Yet the nature, dynamics, and origins of such an essential current have remained elusive. Recently, we found that a compound previously shown to block gating pore currents, 1-(2,4-xylyl)guanidinium (XG), abrogates pacemaking of rodent midbrain dopamine neurons (mDANs) ^7^. Using voltage ramps and pacemaker-clamp, we were able to isolate the tiny current blocked by XG, reminiscent of the elusive *I_pace_*. Since slow pacemaking is such a fundamental behavior for life, we hypothesized that this mechanism is conserved across neuronal and cardiac pacemaker myocytes and species. We therefore investigated whether similar effects of XG would be observed in human mDANs and in rodent sinoatrial node (SAN) pacemaker myocytes and found it to be the case. Dynamic-clamp experiments, in which electrophysiological recordings are coupled to exogenous current injections to either add or subtract conductances, confirmed that the XG-sensitive *I_pace_* is necessary and sufficient for slow pacemaking. Next, we used both a conductance-based model of a mDAN ^17^ and a reduced neuronal model and showed that the voltage dependence of *I_pace_* is ideally suited to be the effector of slow pacemaking. Taken together, our data supports the hypothesis of a conserved mechanism for slow pacemaking, encompassing neuronal and cardiac pacemaker cells.

## Results

### Evidence for a shared mechanism of slow pacemaking across species and systems

We had previously shown that pacemaking of rodent mDANs is blocked by XG and had isolated the current sensitive to XG (**Fig. 1A-C**) ^7^. We first asked whether pacemaking in human mDANs derived from two different induced pluripotent stem cells (hIPSCs) lines (see *Methods*) would be similarly affected by the drug. These neurons were identified as mDANs based on their spontaneous slow firing frequency, hyperpolarization by dopamine and, in some cases, the expression of tyrosine hydroxylase (**Fig. S1**; see *Methods*). Upon superfusion of XG, their spontaneous activity was silenced (2.28 ± 0.44 Hz in control vs 0.12 ± 0.05 in XG, n = 9, p = 0.0039, Wilcoxon test; **Fig. 1D, E**), similarly to what we had observed in rodent models. In particular, as in rodent mDANs, the silencing of the electrical activity by XG does not involve membrane hyperpolarization, contrary to what is seen with dopamine, which is known to open G-protein-coupled inwardly rectifying channels in these cells (**Fig. S1**). Moreover, we observed a current sensitive to XG in human mDANs which is similar to the one observed in rat mDANs (**Fig. 1F**). Taken together, these results suggest that the mechanism of pacemaking is conserved across species in this cell type.

**Fig. 1.**
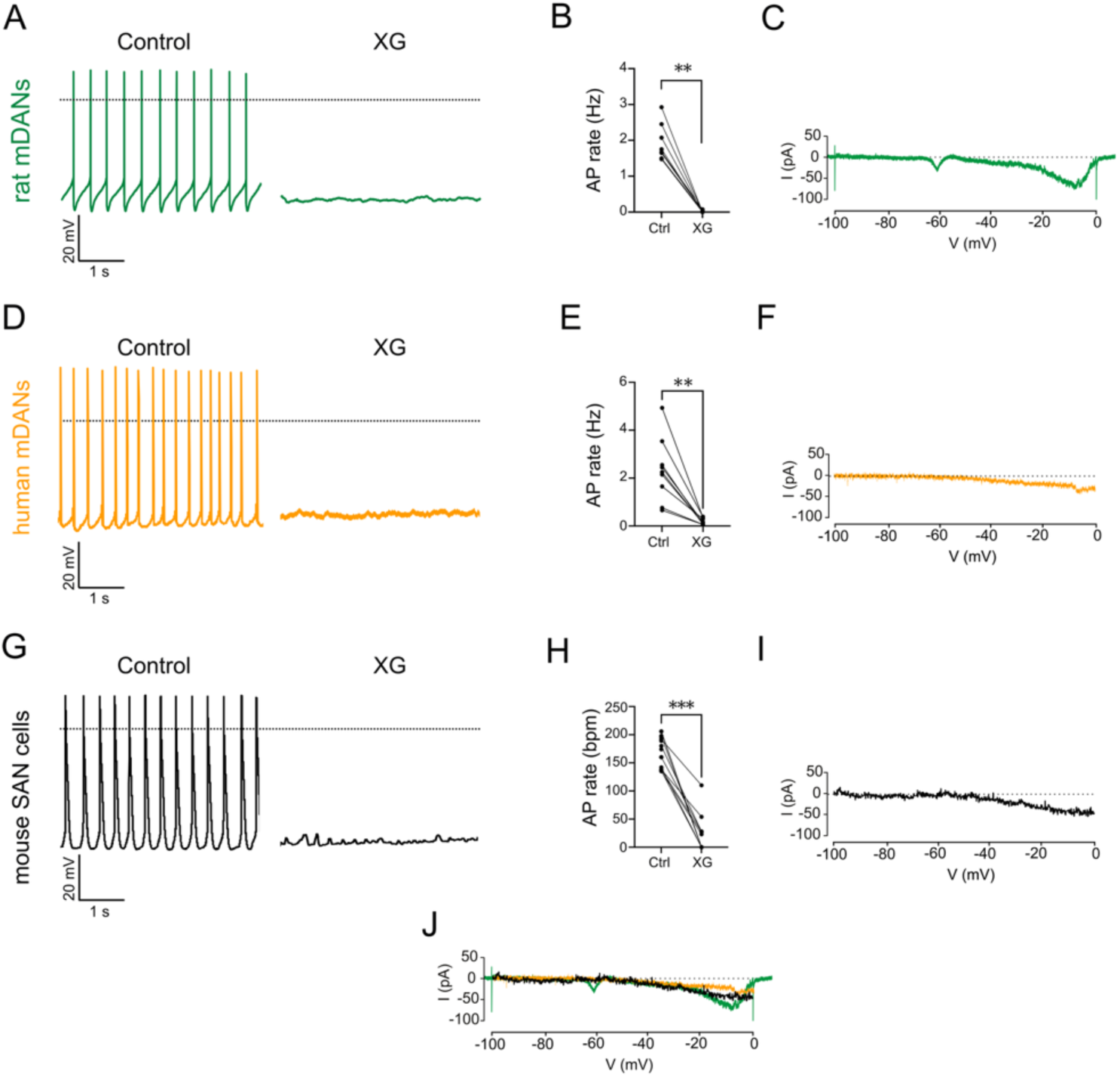
XG inhibits slow pacemaker cells. (**A**) Pacemaking in rat mDANs is inhibited upon (2 mM) XG application. (**B**) Summary data of the effect of the application of 2 mM XG on the firing frequency. XG abolished the firing (1.94 ± 0.18 Hz in control vs 0.02 ± 0.001 Hz in XG, n = 8, p = 0.0078, Wilcoxon test). (**C**) XG-sensitive current isolated from rat mDANs. Inward current at -60 mV is the result from rundown of T-type calcium channels that was observed in one cell. (**A-C**) were adapted from *Jehasse et al. 2021* ^7^. (**D**) Pacemaking in human mDANs is inhibited upon (600 µM) XG application. (**E**) Summary data of the effect of the application of 600 µM XG on the firing frequency (2.28 ± 0.44 Hz in control vs 0.12 ± 0.05 in XG, n = 9, p = 0.0039, Wilcoxon test). (**F**) XG-sensitive current isolated from human mDANs (**G**) Inhibition of pacemaking in mouse SAN myocytes by XG (300 µM) application. (**H**) Summary data of the effect of application of 600 µM XG on the spontaneous pacemaking of SAN myocytes, expressed in beats per minute (bpm). XG strongly reduces pacemaking (166.50 ± 9.11 bpm in control vs 24.55 ± 10.10 bpm in XG, n = 11, p = 0.001, Wilcoxon test). (**I**) XG-sensitive currents isolated from mouse SAN myocytes. (**J**) Overlay of XG-sensitive current from rat (green), human mDANs (orange), and SAN myocytes (black) shows similarities in voltage activation and current amplitude. Dashed lines correspond to 0 mV for voltage traces and 0 pA for current traces.

To investigate whether this finding could be extended to another cell type endowed with pacemaking, we explored the case of SAN pacemaker myocytes of the heart, which are also slow (1-5 Hz), very regular, pacemakers. In addition, similar to mDANs, these cells express high levels of L-type Ca_v_1.3 and HCN channels ^5^. Upon superfusion of XG, we observed a marked decrease of pacemaking in SAN myocytes (from 166.5 ± 9.1 bpm in control to 24.5 ± 10.1 bpm in XG, n = 11, p = 0.001, Wilcoxon test; **Fig. 1G, H**). Slowing of pacemaker activity by XG could not be attributed to interaction with delayed-rectifier K^+^ currents, the L- and T-type Ca^2+^ current (*I_CaL_* and *I_CaT_*), nor with HCN-mediated *I_f_* current (**Fig. S2**), but to inhibition of a tiny inward “pacemaker” current that we refer to as *I_pace._* In SAN myocytes, *I_pace_* showed similar characteristics to those previously observed in mDANs (**Fig. 1I, J**), including activation at negative voltages (-50 mV), voltage-dependent current increase along the diastolic depolarization voltage range (corresponding to the neuronal interspike) and peaking at ∼-20 mV, a voltage more depolarized than the threshold of the action potential in both cell types. Moreover, XG inhibited the spontaneous activity of isolated SAN pacemaker myocytes with apparent concentration for half-inhibition (IC_50_) of 32 µM, smaller than what we previously observed in rodent mDANs from acute brain slices ^7^. The Hill coefficient was 2.5 (**Fig. S2D**). Consistently, XG decreased the spontaneous pacemaking rate of isolated Langendorff-perfused hearts (**Fig. S2E**). Taken together, these data indicate XG-sensitive *I_pace_* as a shared mechanism for slow pacemaking in mDANs and in SAN pacemaker myocytes.

### Modeling the pacemaker current confirms its ability to sustain slow pacemaking

We used a well-established conductance-based model to study slow pacemaking (**Fig. S3A, B**) ^17^. Since this model is set to pace steadily without a pacemaker current, it relies on shifting the activation curve of Na^+^ (*g_Na_*) and L-type Ca^2+^ (*g_CaL_*) conductances to artificially hyperpolarized potentials. We adjusted these activation curves to their measured physiological values in mDANs ^18,19^ (**Fig. S3C**), which led to a loss of spontaneous firing and a shift from type I to type II excitability (**Fig. S3B**). We then modeled the pacemaker conductance (*g_pace_*) from the current-to-voltage curve of the XG-sensitive *I_pace_* (**Fig. S3D**) and implemented it into the adjusted model (**Fig. S3E**). We were able to recover pacemaking activity with a voltage range of activation of *g_pace_* similar to what had been observed experimentally (**Fig. 2A, B**) ^7^, as well as to recover type I excitability **(Fig. S3B**). Currentscape analysis ^20^ revealed the crucial contribution of *g_pace_* during pacemaking (**Fig. 2A**). We then measured the total current between two spikes and observed that the resulting current is driven by *g_pace_* (**Fig. 2B**) and has a small amplitude similar to previous experimental observations in mDANs ^21^. Finally, we measured the charge transfer of each conductance during an interspike interval and a full cycle (**Fig. 2C**). Interestingly, the charge carried by *g_pace_* (-191 fC) is much smaller than the charge carried by other conductances such as *g_CaL_* (-1458 fC) or H-type conductance (*g_H_*) (-2077 fC) during the interspike interval. Overall, these data highlight the essential role of the small conductance *g_pace_* in slow pacemaking.

**Fig. 2.**
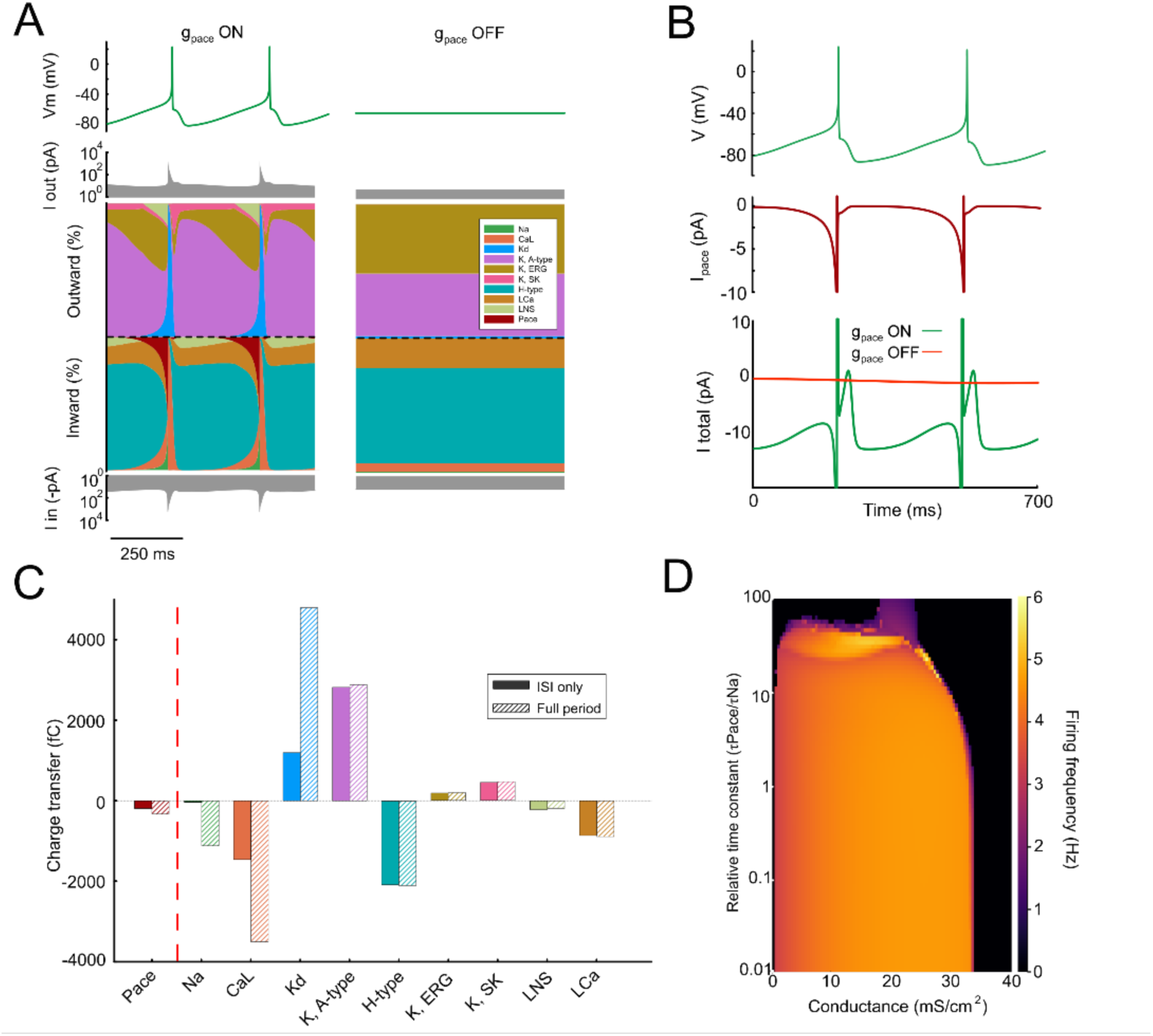
Fast activating pacemaker conductance allows slow pacemaking in a mDAN conductance-based model. (**A**) Currentscape analysis reveals that, although its contribution seems small in comparison to other conductances, *g_pace_* is crucial to generate slow pacemaking. (**B**) Simulated pacemaking activity in our adjusted pacemaking model (green) with *I_pace_* dynamic below in red, showing a small amplitude during the interspike interval as previously observed ^7^. Bottom panel corresponds to total current amplitude from our adjusted model with (green) and without (red) the addition of *g_pace_*. The total current observed is similar to what has been calculated ^13^. (**C**) Total charge transfer of each conductance during the interspike interval (ISI) and the full period of the simulation. Charge transfer was calculated by measuring the area under the curve in (**A**). (**D**) Heatmap analysis showing the multiplicative time constant of *g_pace_* activation relative to *g_Na_* according to the conductance density in which our adjusted model generates slow pacemaking.

We next investigated the dynamical range of *g_pace_* which is necessary to maintain a steady low frequency spontaneous activity (**Fig. 2D**). We explored both the density and activation time constant of *g_pace_*, relative to the activation of Na_v_ channels, and found that the model remains stable as long as the activation of *g_pace_* is very fast, close to instantaneous. Moreover, *g_pace_* can have a wide range of densities, making the phenomenon robust. To verify that these observations are not limited to the particular parameters used, we performed the same analysis using random parameters in which slow pacemaking can be observed (**Fig. S4**). In all cases, we observed similar requirements for the parameters of *g_pace_*. Moreover, the addition of *g_pace_* into the model increased ion channel degeneracy (**Fig. S5**), especially for *g_Na_* and *g_CaL_*, suggesting that it may stabilize neuronal output in the face of biological variability ^22–25^.

To further explore the structure of this increased degeneracy, we performed principal component analysis on the pacemaking solutions of the adjusted model. The analysis revealed no preferred direction of variability, indicating that the contribution of *g_pace_* does not impose a dominant constraint on the system but instead distributes flexibility across all conductances (**Fig. S5C**). This isotropic degeneracy suggests that slow pacemaking remains robust even when the underlying ion channel expression varies extensively, emphasizing the stabilizing role of *g_pace_* in maintaining reliable activity.

### Dynamic clamp experiments confirm that g_pace_ is necessary and sufficient for slow pacemaking

Next, we used the dynamic-clamp approach to test how *g_pace_* regulates slow pacemaking in spontaneously active mDANs from *substantia nigra pars compacta* (*SNc*) recorded *ex vivo* (**Fig. 3A**). First, we injected several values of negative conductance and evaluated the percentage of decrease in firing frequency (**Fig. 3B**). We observed a frequency decrease of 50% at -0.06 nS and complete silencing at -0.3 nS (1.54 Hz in control vs. 0 Hz with *g_pace_* = - 0.3 nS, n = 20, N = 3, p = <0.0001, Wilcoxon test; **Fig 3B**). Injecting positive values for *g_pace_* increased the firing frequency as expected, leading to a depolarization block at the highest value tested (above 0.3 nS, **Fig. 3C**). Finally, we preincubated *SNc* DANs with XG to silence their activity and injected positive *g_pace_* to test whether we could rescue slow pacemaking (**Fig. 3D**). Overall, the rescue was successful in all trials (n = 11) at a variable intensity of *g_pace_* which is to be expected because of the diversity of the biophysical properties of mDANs ^26^. Finally, we investigated the kinetics of *g_pace_*. mDAN pacemaking was first silenced with XG and we rescued the activity upon *g_pace_* injection (**Fig.4A, upper panel**). However, when changing the activation time from 1 ms to 30 ms at a constant conductance value, we observed that mDANs are more likely to generate a depolarization block with slower dynamics (**Fig. 4A-C**). This is entirely consistent with our model (**Fig. 4D**). The latter experiment demonstrates that our modeling and experimental data are congruent and confirm that a fast and tiny current is required for slow pacemaking.

**Fig. 3.**
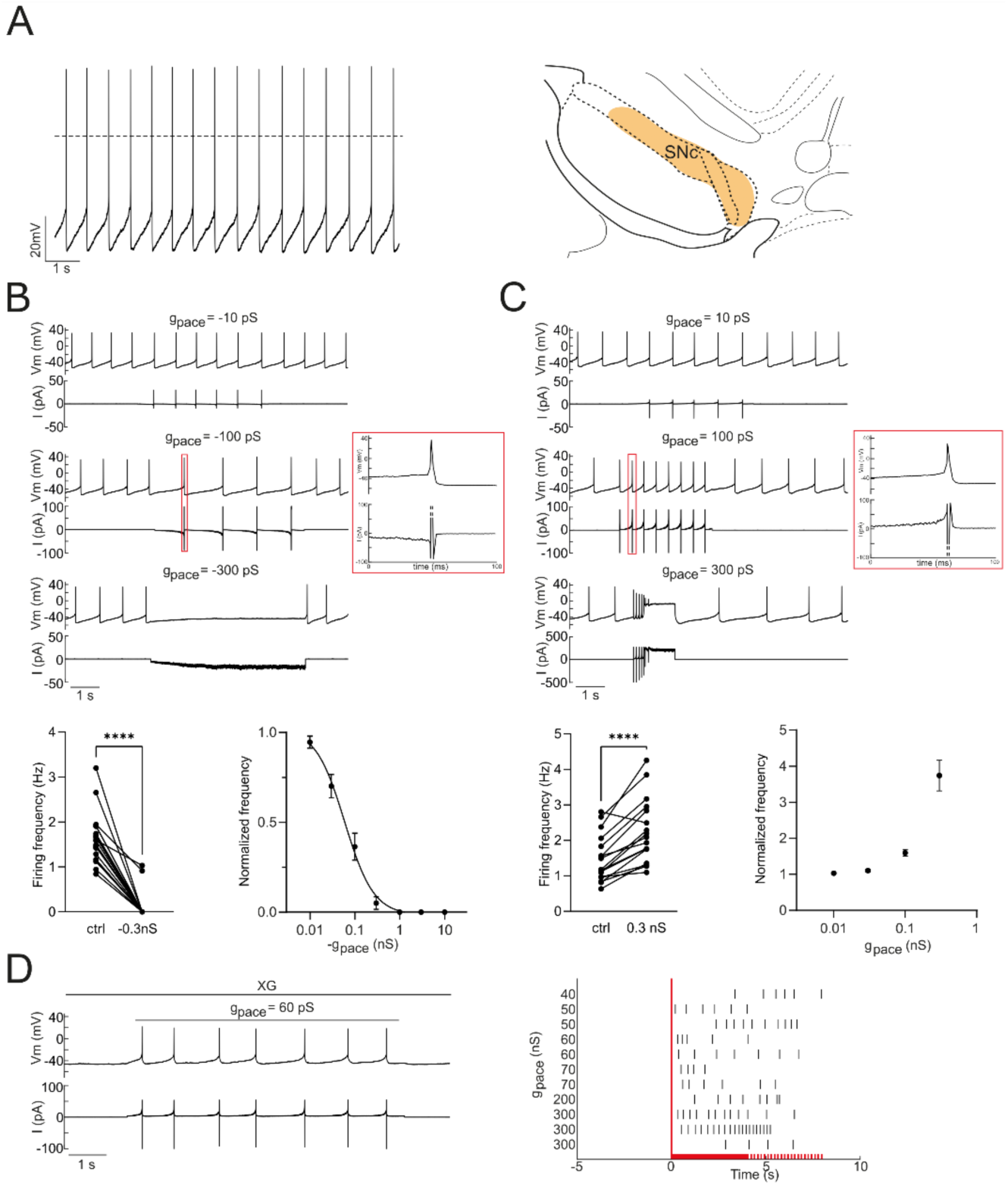
The XG-sensitive conductance (*g_pace_*) modulates pacemaking in *SNc* DANs. (**A**) *Ex vivo* pacemaking in a mouse *SNc* (orange shade) DAN. (**B**) Silencing of a *SNc* DAN upon injection of negative *g_pace_* (upper panel). Summary data of the effect of negative *g_pace_* on the firing frequency (lower panel). Negative *g_pace_* abolished pacemaking at -0.3 nS (1,54 Hz (interquartile range: 1,193 - 1,864 Hz) in ctrl vs. 0 Hz (interquartile range: 0) with *g_pace_* = - 0,3 nS, n = 20, N = 3, p = <0,0001, Wilcoxon test). (**C**) Excitation and overexcitation (depolarization block) of a *SNc* DAN upon positive *g_pace_* (upper panel). Summary data of the effect of positive *g_pace_* on the firing frequency (lower panel). *g_pace_* increased the firing frequency (1,48 ± 0,66 Hz in ctrl vs. 2,26 ± 0,91 Hz with *g_pace_* = 0,3 nS, n = 17, N = 3, p = <0,0001, paired t test). (**D**) Rescue of pacemaking after preincubation with (2 mM) XG and complete cessation of pacemaking. Firing pattern and *g_pace_* value upon rescue display broad large variability (right). Time frame of steady state *g_pace_* (bold red line). Time frame of manual deactivation of *g_pace_* (discontinuous red line).

**Fig. 4.**
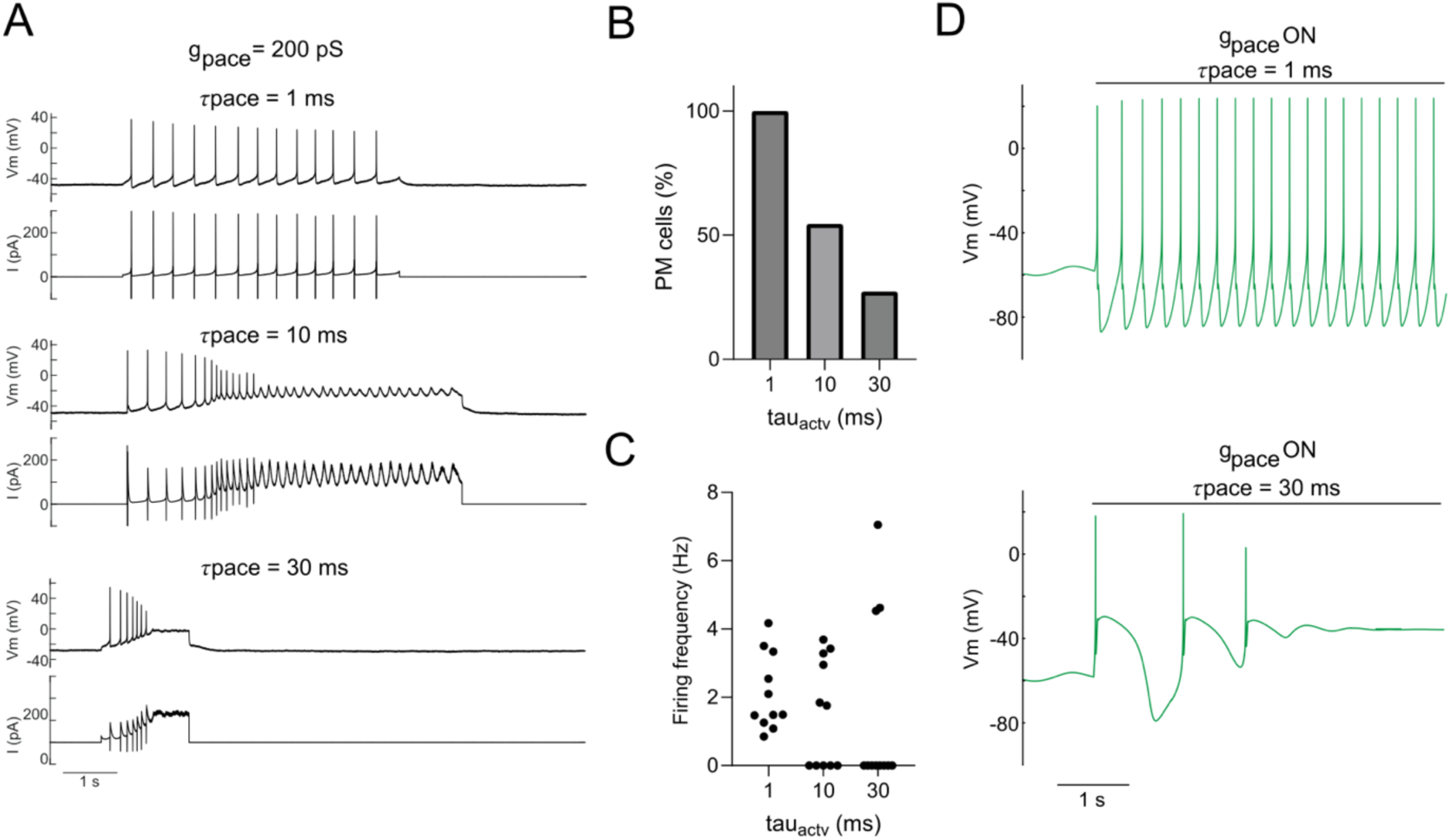
The XG-sensitive conductance (*g_pace_*) needs to be fast to obtain slow pacemaking. (**A**) Rescued pacemaking from mouse *SNc* DANs upon 2 mM XG superfusion. Increasing activation time of *g_pace_* (1_pace_) generates a depolarization block. (**B**) Percentage of pacemaking cells decreases with slower activation time (n=11 for each group). (**C**) The amount of slow firing cells decreases with slower activation time (n=11 for each group). (**D**) Increased t_pace_ in our model generates a depolarization block, similarly to what we observed *in vitro*.

### Respective roles of I_pace_ and other conductances for slow, regular pacemaking

We next sought to understand how important *g_pace_* is for slow pacemaking. We reduced our conductance-based model to a minimal one, keeping only *g_Na_* and *g_Kd_* for spike generation, as well as *g_LNS_* (leak conductance) to set the membrane at the desired potential. Adding *g_pace_* in this minimal configuration enabled slow pacemaking similarly to our complete model (**Fig. 5A**). In this model, slow pacemaking is still maintained with fast activating *g_pace_* over a wide range of densities, from 4.5 mS/mm² to 35.5 mS/mm² (**Fig. 5B**). Since our data showed that this minimal configuration is sufficient to create slow pacemaking, we investigated the role of the other conductances, especially A-type K^+^ (*g_KA_*), H-type (*g_H_*) and Ca^2+^-dependent K^+^ (*g_KSK_*) conductances. These have been shown in slices to help maintaining the firing frequency and regularity constant during membrane potential changes caused by excitatory and inhibitory inputs ^3,21,27^. We introduced current noise in our minimal model and observed that the pacemaking becomes irregular (**Fig. 5C**). In our complete model, the firing frequency is robust to current injection, similarly to what has been observed by others ^3,21,27^. Moreover, we also observed that these conductances, and especially *g_KSK_*, help to reduce the slope of the firing frequency – injected current relationship, which is expected (**Fig. 5D**).

**Fig. 5.**
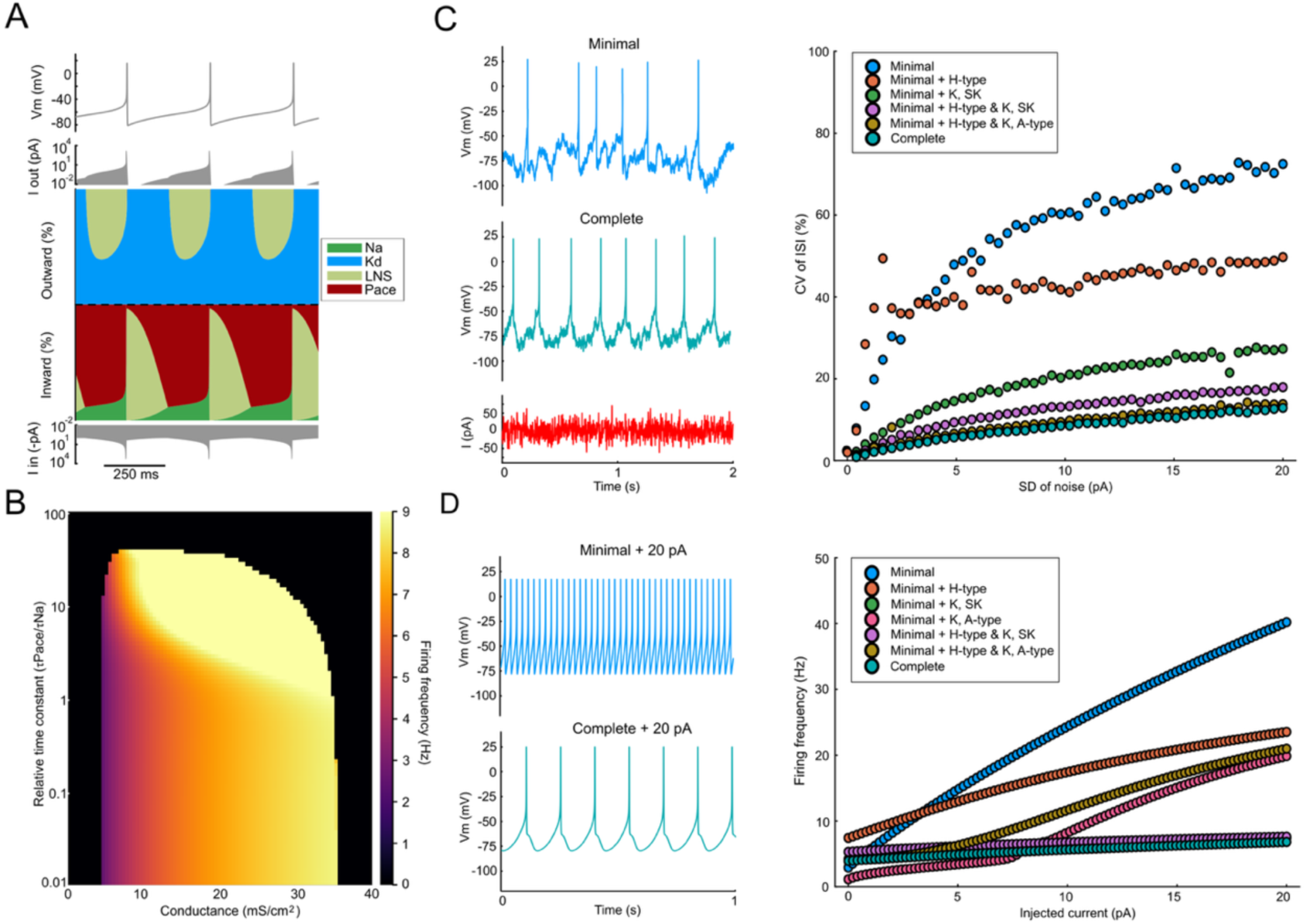
Assessment of the conductances contribution in slow pacemaking. (**A**) Currentscape analysis from a minimal model including *g_Na_*, *g_Kd_*, *g_LNS_* and *g_pace_* shows that it is possible to generate slow pacemaking, similarly to what we observed *ex vivo* in the absence of other conductances ^7^, and that *g_pace_* contribution is maximal during the interspike interval. (**B**) Heatmap analysis showing the multiplicative time constant of *g_pace_* activation relative to *g_Na_* according to the conductance density in which our minimal model generates slow pacemaking. (**C**) Robustness of slow pacemaking in response to noise (in red). In the minimal model (blue), current noise leads to irregular spiking, while the complete model (teal), current noise injection does not alter pacemaking regularity (left panel). Right panel corresponds to the effect of increasing the standard deviation (SD) of noise current to the coefficient of variation (CV) of the interspike interval (ISI). The addition of *g_H_*, *g_KSK_* and *g_KA_* increases the robustness of slow pacemaking. In order to activate *g_KSK_*, *g_LCa_* was added. (**D**) Effect of the other conductances on the firing frequency upon continuous current injection. Left panel shows example of the firing frequency of the minimal (blue) and complete (teal) upon current injection of 20 pA. Right panel shows the effect of the combination of several conductances on the FI relationship. The results related to *g_KSK_* addition only are located behind the complete and combination of *g_KSK_* and *g_H_* data points.

We then challenged the requirement of a voltage-gated activation for *g_pace_* in order to obtain slow pacemaking. We replaced *g_pace_* by a linear leak conductance similar to NALCN channels (*g_Nalcn_*), as they had been suggested by some authors to be the main drivers of pacemaking in some mDANs ^13,28–30^. *g_Nalcn_* failed to sustain slow pacemaking (**Fig. S6**). In fact, low or high density of *g_Nalcn_* could not generate spontaneous firing (**Fig. S6B**), while activity could be observed in a small window of density, going from type II excitability to fast pacemaking type I excitability (**Fig. S6C**). Our data therefore supports the notion that slow pacemaking requires a current with a specific voltage-dependent activation during the interspike interval, not a linear one.

## Discussion

Slow pacemaking is a unique property which is observed in a subset of cells that play a major role in rhythmic biological processes. However, the precise mechanism(s) underlying slow pacemaking is (are) still controversial. Recently, we found that XG inhibits the pacemaking of rodent mDANs and were able to isolate the current blocked by this compound ^7^. Here we show that this effect can be extended to human mDANs and to mouse SAN pacemaker myocytes. Remarkably, the XG-sensitive conductance or *g_pace_* has very similar voltage-dependence in mDANS and SAN myocytes, starting to operate at ∼-50 mV and slowly increasing in amplitude towards more depolarized potentials. In our minimal conductance-based model, we show that such a voltage dependence is able to generate slow pacemaking, whereas addition of a linear conductance such as NALCN is unable to induce this firing pattern.

In our adjusted conductance-based model of a mDAN, we show that addition of *g_pace_* is able to induce biologically realistic pacemaking. The properties of *g_pace_* allow to extend ion channel degeneracy, which makes our model more robust to biological variability (**Fig. S5**). In this work, we describe key properties that *I_pace_* should have. First, its activation should be fast. Second, its amplitude can vary within a relatively large range, which guarantees robustness. In addition, we show that, while large membrane conductances, such as *g_CaL_*, *g_KA_* or *g_H_* are not required for slow pacemaking ^3,7^, *I_pace_* is sufficient to generate slow automaticity in both the minimal and complete models.

Dynamic-clamp experiments confirmed the importance of *g_pace_* (**Fig. 3**). Moreover, the range of conductances to either rescue slow pacemaking upon XG perfusion or to silence neuronal activity was a small fraction of the cell conductance, around 3 nS in rodent mDANs ^7^. Overall, setting *g_pace_* to 0.3 nS was enough in both the negative conductance and rescue experiments, confirming the fact that pacemaking relies on a tiny current. These data support our hypothesis that the pacemaker channel should have a unitary conductance value in the fS range ^7,14^. In fact, if *g_pace_* were to have a single channel conductance similar to the main pore of Na_v_ channels (around 20 pS), it would imply that slow pacemaking is sustained by at most 15 channels. Because of intrinsic stochasticity of voltage-dependent channel gating, this would lead to very irregular firing in neurons and in the heart.

In mDANs, the role of several ion channels in pacemaking has been investigated over the years. L-type Ca_v_ channels were considered, since their pharmacological blockade with high concentrations of dihydropyridines silenced these neurons ^31–33^. Further studies employing lower, pharmacologically relevant concentrations of L-type blockers did not confirm these previous data ^7,8,34,35^, therefore suggesting an off-target effect of dihydropyridines at high concentrations. This is consistent with our numerical simulations in the present study, since our minimal model predicts pacemaking without a Ca_v_ conductance. L-type Ca_v_1.3 channels are now considered as linear amplifier of the firing frequency in mDANs ^8^. NALCN channels were also considered as a pacemaker mechanism, as they are expressed in most rhythmic neurons ^36^. However, further studies showed controversial results, in which NALCN gene deletion or pharmacological blockade suppressed ^28–30,37^ or did not impact pacemaking activity ^7,29^. In this regard, our simulations show that a linear depolarizing conductance is unlikely to sustain stable pacemaking.

SAN pacemaking has been linked to either voltage-dependent activation of HCN channels, or to a process of spontaneous cyclical Ca^2+^ release by cardiac ryanodine receptors coupled to the Na^+^/ Ca^2+^ exchanger (a process referred to as the “Ca^2+^ clock” ^38^). However, genetic silencing of cardiac HCN channels leads to bradycardia and arrhythmia ^39^, without arresting heart pacemaking. In addition to HCN channels and the Ca^2+^ clock, L-type Ca_v_1.3 channels ^40^ and T-type Cav3.1 ^41^ channels have been proposed to provide inward current in the diastolic phase (corresponding to neuronal interspike interval) to generate pacemaking. In this regard, further work indicated also that Ca_v_1.3 channels (i) constitute the main route for entry of Ca^2+^ in diastolic phase, (ii) activate the Ca^2+^ clock ^42^ and (iii) mediate up-regulation of pacemaking by catecholamines with HCN channels ^12^. However, concomitant genetic ablation of Ca_v_1.3 and Ca_v_3.1 channels strongly reduces (∼60%), but still does not abrogate SAN pacemaking, even under conditions of HCN and Nav channel inhibition ^11^. These data strongly suggest the existence of an additional ionic mechanism supporting slow cardiac pacemaking and accounting for persistence of automaticity when other ion channels involved in the generation of diastolic depolarization are inhibited. The properties of *I_pace_* in DA neurons, including voltage dependency, insensitivity to gadolinium ^7^ and to inhibition of the Na^+^/Ca^2+^ exchanger, suggest that in the SAN, *g_pace_* is not generated by TRPM4 ^43^ or TRPC ^44^ channels, but rather accounts for a previously overlooked ionic mechanism of pacemaking.

In conclusion, our study unravels a new ionic current underlying spontaneous electrical activity, shines new light on the controversial mechanism of slow pacemaking in excitable cells and indicates a commonality across systems, an observation that is very novel and surprising.

## Materials and Methods

### Modeling

#### Programming language

The Julia programming language was used in this work ^45^. Numerical integration was realized using *DifferentialEquations.jl*. Fitting was realized using *LsqFit.jl*. PCA was realized using *MultivariateStats.jl*.

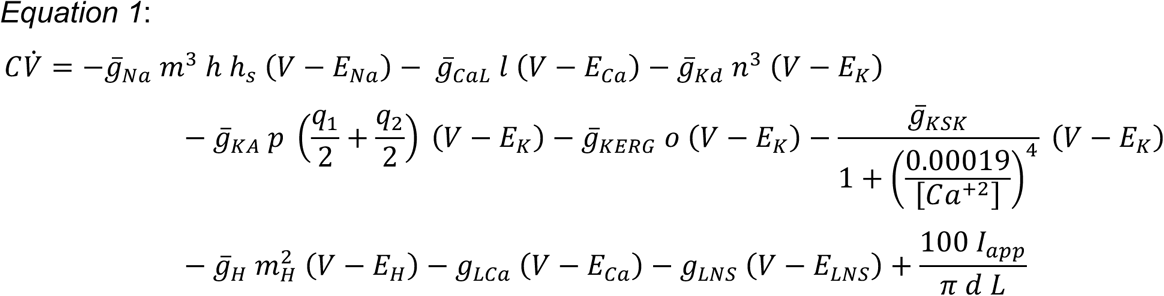

Here, the dot notation represents time derivative (in /ms), V is the membrane potential (in mV), and C the membrane capacitance (in µF/cm²). The ionic currents are: *I_Na_* (transient sodium), *I_CaL_* (L-type calcium), *I_Kd_* (delayed rectifier potassium), *I_KA_* (A-type potassium), *I_KERG_* (ether-à-go-go potassium), *I_KSK_* (SK potassium), *I_H_* (hyperpolarization-activated current), *I_LCa_* (calcium leak), and *I_LNS_* (non-specific leak). Reversal potentials are denoted by E. The intracellular calcium concentration [Ca^+2^] follows voltage-dependent kinetics as in (*13*). The soma geometry is defined by its diameter d and length L (in µm). The applied current is *I_app_* (in pA).

Maximal conductances are denoted by the bar notation (in mS/cm²), representing the channel conductance when fully open. Gating variables follow first-order voltage-dependent kinetics of the following form.

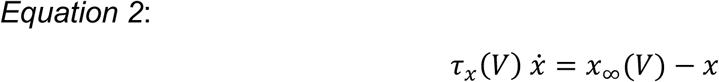

Here, x is the gating variable, τ_x_(V) its voltage-dependent time constant, x_∞_(V) its steady-state activation or inactivation function. Variables that were adjusted in this work are specified in the following section.

Unless otherwise stated, the original model parameters were used (Table S1) (*13*).

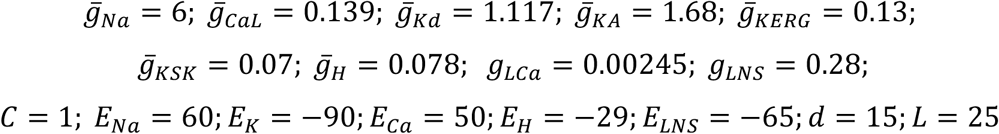

#### Adjustment of the mDAN conductance-based model and fitting of the pacemaking current

Before introducing the pacemaking current, activation of the transient sodium and L-type calcium channel was realized. In the original model (*13*), the steady-state activation functions were defined as follows.

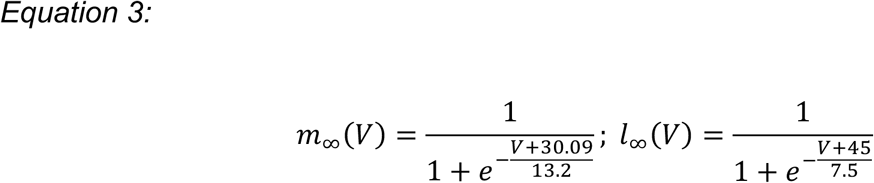

These sigmoidal functions have half-activation voltages of −30.09 mV and −45 mV, respectively, which are substantially more hyperpolarized than values reported experimentally. In the adjusted model, both activation curves were shifted to −10 mV. The ultraslow inactivation of the transient sodium current (h_s_) was disabled, as it was an adjustment to generate slow pacemaking in the original model ^17^.

The pacemaking current was fitted to a previously published voltage–current relationship (*3*) (**Fig. 1C**). The pacemaking current was modeled as a non-inactivating current, which provided an accurate fit without unrealistic parameter estimates.

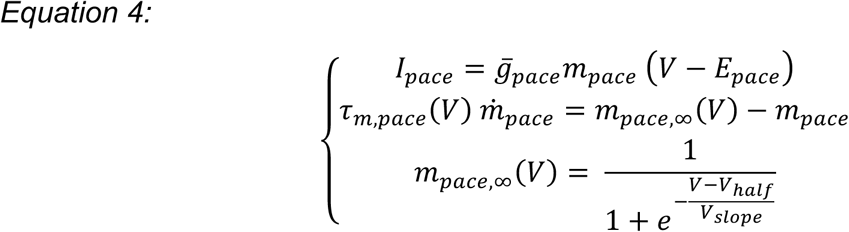

Because the fit was performed on steady-state voltage–current data, no biological information was available on the time constant of pacemaking activation. We therefore set

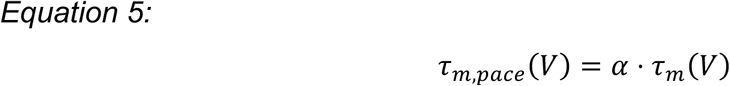

where τ_m_(V) is the activation time constant of the transient sodium current.

Fitting yielded the following parameter values, given to one decimal place.

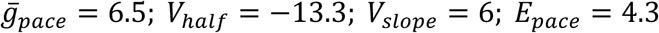

Since this pacemaking conductance corresponds to a specific neuron, it was varied throughout the study to test robustness rather than fixed at the fitted value. Unless otherwise stated, the following parameters were used in the adjusted model.

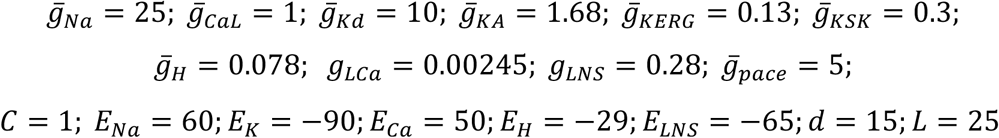

#### Modeling of a NALCN current

In **Fig. S6**, the NALCN conductance was modeled as a voltage-independent sodium current.

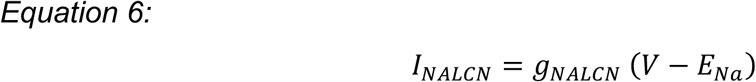

#### Degenerate population building

In **Fig. S5**, two degenerate populations (N=200) exhibiting pacemaking were generated for the original and adjusted models. The procedure was identical for both populations. All ionic conductances were randomly assigned using a uniform distribution between 0 and an upper bound value, corresponding to the ordinates of **Fig. S5B**. Each model was simulated and retained in the population if its firing frequency was between 1 and 5 Hz, if the spike peak exceeded 0 mV, and if the minimum membrane potential was between −90 and −60 mV.

### Cell Culture

#### Cell lines

Two human induced pluripotent stem cell (hiPSC) lines were used in this study: GM23338 (gift from Prof. Ira Mercedes Espuny Camacho, GIGA Neurosciences, University of Liège) and a human TH-reporter iPSC line expressing mCherry (gift from Dr. Aleksandar Rakovic, Institute of Neurogenetics, University of Lübeck) ^46^. Prior to differentiation, cells were maintained for 2–3 consecutive passages in E8 medium using standard EDTA-based passaging when confluency reached 70–80%.

#### Differentiation to midbrain dopaminergic neurons

Differentiation was carried out using a slightly modified version of the protocol published by Stathakos *et al*. (2019) ^47^. Briefly, hiPSCs were cultured in E8 medium for up to 4 days before switching to neural induction medium consisting of N2B27 supplemented with 100 nM LDN193189, 10 μM SB431542, 300 ng/mL SHH, and 0.6 μM CHIR99021.

This was considered neuralization day 0. Cells were passaged on polyornithine/laminin-coated plates at days 3 and 7 using Accutase at a density of 4 × 10^4^ cells/cm² in induction medium with 10 μM Y27632 or 1:100 RevitaCell.

From day 11, the medium was replaced with progenitor expansion medium (N2B27 with 20 ng/mL BDNF, 20 ng/mL GDNF, 0.2 mM ascorbic acid, and 10 μM Y27632). Upon confluence, cells were passaged every 3-4 days in this medium. Terminal differentiation started on day 16 by seeding cells at 5,000 cells/cm² onto PO/laminin-coated coverslips in N2B27 supplemented with 20 ng/mL BDNF, 20 ng/mL GDNF, 0.2 mM ascorbic acid, 500 μM db-cAMP, 10 μM DAPT, and 10 μM Y27632. Cells were allowed to attach for ∼1 hour before flooding wells with the same medium. Media was refreshed every 3 days with 50% volume exchange.

### Tissue preparation

#### Animals

For dynamic-clamp experiments, adult male C57BL/6N (Charles River Laboratories) mice (8 to 14 weeks of age) were used. A maximum of 5 mice were housed per cage and single animal housing was avoided when possible. Animals were maintained on a 12-hour light-dark cycle and provided with water and food ad libitum. All experimental procedures involving mice were approved by the German Regional Council of Darmstadt.

For SAN cells experiments, the study is in accordance with the Guide for the Care and Use of Laboratory Animals published by the US National Institute of Health (NIH Publication No. 85-23, revised 1996) and European directives (2010/63/EU). All mice used were from the C57BL/6J background (Charles River and Janvier labs).

Experimental procedures were approved by the Ethical Panel of the University of Montpellier and the French Ministry of Agriculture (protocol no: 2017010310594939). Animals were housed in the IGF animal facility with free access to food and water and were exposed to 12 h light/dark cycles in a thermostatically controlled room (21–22 °C) with 40−60% humidity.

#### Slice preparation for mouse midbrain dopamine neurons

Animals were anesthetized by intraperitoneal injection of ketamine (250 mg/kg; Ketaset, Zoetis) and medetomidine hydrochloride (2.5 mg/kg; Domitor, OrionPharma) before intracardial perfusion using ice-cold ACSF consisting of the following: 50 mM sucrose, 125 mM NaCl, 2.5 mM KCl, 25 mM NaHCO_3_, 1.25 mM NaH_2_PO_4_, 2.5 mM glucose, 6 mM MgCl_2_, 0.1 mM CaCl_2_, and 2.96 mM kynurenic acid (Sigma-Aldrich), oxygenated with 95% O_2_ and 5% CO_2_. Rostral coronal midbrain slices (bregma, −2.92 to −3.16 mm) were sectioned into 250-µm slices using a vibrating blade microtome (VT1200s, Leica). Before the experiment, slices were kept at 37°C for 1 hour in oxygenated extracellular solution containing the following: 22.5 mM sucrose, 125 mM NaCl, 3.5 mM KCl, 25 mM NaHCO_3_, 1.25 mM NaH_2_PO_4_, 2.5 mM glucose, 1.2 mM MgCl_2_, and 1.2 mM CaCl_2_.

#### Isolation of mouse sinoatrial node myocytes

Sinoatrial node myocytes were isolated as previously described ^48^. Briefly, Sinoatrial tissue (SAN) strips were immersed into “low Ca^2+^” Tyrode’s solution containing (in mM): 140 NaCl, 5.4 KCl, 0.5 MgCl_2_, 1.2 KH_2_PO_4_, 50 taurine, 5.5 D-glucose, 1 mg/mL BSA, and 5 Hepes (adjusted to pH 6.9 with NaOH) for 5 min. The tissue was then transferred into the low-Ca^2+^ solution containing Liberase TH (0.15 mg/mL, Roche) and Elastase (0.48 mg/mL, Worthington). Digestion was carried out for 15-20 min at 36 °C. SAN strips were then washed in “Kraftbrühe” (KB) solution containing 100 L-glutamic acid potassium salt, 10 L-Aspartic acid potassium salt, 25 KCl, 80 KOH, 10 KH_2_PO_4_, 2 MgSO_4_, 5 Creatine, 0.5 EGTA, 20 D-glucose, 20 taurine, 1 mg/mL BSA, and 5 Hepes (adjusted to pH 7.2 with KOH). Single myocytes were isolated from the SAN tissue by manual agitation using a flame-forged Pasteur’s pipette in KB solution at 36 °C. For recovering of pacemaker activity, Ca^2+^ was gradually reintroduced into the cells’ storage solution to a final concentration of 1.8 mM. Cells were then stored at room temperature until use.

### Electrophysiological Recordings

#### Midbrain dopamine neurons

For rat mDANs, the recordings were obtained from previous work ^7^

For human mDANs, electrophysiological properties of differentiated neurons were assessed using both on-cell and whole-cell patch-clamp recordings performed at 30°C with a Multiclamp 700B amplifier and Clampex 11.1 software. The extracellular solution contained (in mM): 140 NaCl, 5 KCl, 2 CaCl_2_, 2 MgCl_2_, 15 HEPES, and 10 D-glucose; pH was adjusted to 7.3 with NaOH. The intracellular pipette (3-5 MΩ) solution contained (in mM): 135 K-gluconate, 10 NaCl, 0.5 CaCl_2_, 15 HEPES, 2 Na_2_ATP, and 5 EGTA; pH adjusted to 7.3 with KOH, osmolarity ∼ 300 mosm. Putative *substantia nigra pars compacta* (*SNc*)-type mDANs were identified by similar characteristic electrophysiological features we used to identify rodent mDANs ^7,49–51^, including spontaneous pacemaking activity (0.5–5 Hz), action potential duration > 1.35 ms (measured at half-amplitude), a prominent hyperpolarization-activated inward current (*I_H_*), and membrane hyperpolarization > 5 mV in response to 100 μM dopamine from a holding potential of -60 mV. *I_H_* was observed in many, but not all neurons satisfying these criteria. To test the inhibition of pacemaker activity, 600 μM of 1-(2,4-xylyl)guanidinium mesylate (XG) was applied to the bath solution. XG was produced as previously described ^7^.

For dynamic-clamp recordings in mouse *SNc* DANs, slices were placed in a heated recording chamber (37°C) that was superfused with the oxygenated extracellular solution with a flow rate of 2 to 4 ml/min. CNQX (20 µM), DL-AP5 (10 µM), gabazine (4 µM; SR95531), and were added to inhibit respectively excitatory and inhibitory synaptic transmission. Neurons were visualized using infrared differential interference contrast video microscopy with a digital camera (VX55, Till Photonics) connected to an upright microscope (Axioskop 2, FSplus, Zeiss). Patch pipettes were pulled from borosilicate glass (GC150TF-10, Harvard Apparatus, Holliston, MA, USA) using a temperature controlled, horizontal pipette puller (DMZ-Universal Puller, Zeitz). Patch pipettes (3 to 5 MΩ) were filled with a solution containing the following: 135 mM K-gluconate, 5 mM KCl, 10 mM HEPES, 0.1 mM EGTA, 5 mM MgCl_2_, 0.075 mM CaCl_2_, 5 mM Na adenosine 5′-triphosphate, 1 mM Li guanosine 5′-triphosphate, and 0.1% Neurobiotin, adjusted to a pH 7.35 with KOH. For pharmacology, slices were preincubated for at least 10 min with XG mesylate (2 mM) before recording. Recordings were performed using an EPC-10 patch-clamp amplifier (HEKA Elektronik) with a sampling rate of 20 kHz and a low-pass filter (Bessel, 5 kHz). For analysis, recordings were further digitally filtered at 1 KHz. A real-time Linux-based data acquisition program (RTXI; http://rtxi.org) was used to inject online calculated currents (acquisition rates of 10 kHz) of the modelled XG-sensitive conductance according to the following equation:

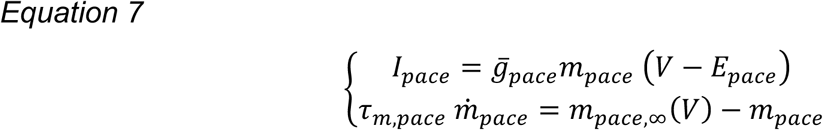

V is the membrane voltage in mV, E_pace_ the reversal potential in mV and *g_pace_* is the conductance in nS, according to the fit in **Fig.S3D**.

#### Heart rate recordings in Langendorff-perfused mouse hearts

We performed ECG recordings in isolated Langendorff-perfused mouse hearts as previously described ^52^. Mice were deeply anesthetized by intraperitoneal injection of 0.3 mL of physiological NaCl solution containing Ketamine (0.1 mg/g, Imalgène) and Xylazine (0.01mg/g, Rompun 2%, Bayer), followed by a second injection of 0.3 mL of solution containing Pentobarbital (150 µL Euthasol / 10 mL NaCl physiological solution). 0.5 mL of NaCl solution containing Heparin (25000 U.I) was injected to avoid the formation of intracardiac blood clots. Hearts were removed via thoracotomy following negativity of the tail sensitivity test. Hearts were cannulated with an aortic cannula, mounted on a Langendorff apparatus. The heart was set in a recording chamber (Warner Instruments, Harvard Apparatus, Holliston, Massachusetts, USA). During recordings, the temperature was maintained at 36.5-37 °C by a two-way, bath and perfusion pipette temperature controller (model TC-344B, Warner Instruments). The ECG was continuously recorded with bipolar electrodes (model SNE 100, Science Products, Hofheim, Germany) placed at the right atrium and apex, connected to a recording amplifier (ADInstruments, Colorado Springs, Colorado, USA).

#### Mouse sinoatrial node myocytes

SAN cells were harvested in custom-made chambers with glass bottoms for cell attachment and superfused with Tyrode solution containing (in mM): 140 NaCl, 5.4 KCl, 1 MgCl_2_, 1.8 CaCl_2_, 5.5 D-glucose and 5 Hepes (adjusted to pH 7.4 with NaOH). warmed at 36°C before recording.

XG-sensitive, delayed-rectifier potassium (*I_K,r_*) and “funny” (*I_F_*) currents were recorded in extracellular Tyrode’s solution under the standard whole-cell configuration whereas action potentials were measured by perforated patch clamp technique with escin (0.04 mg/mL, Sigma-Aldrich) also in extracellular Tyrode’s solution. For *I_F_* recordings, 2 mM BaCl_2_ was added to Tyrode’s solution to block *I_K1_*. Electrodes had a resistance of 3–4 MΩ when filled with a solution containing (in mM): 80 K-aspartate, 50 KCl, 1 MgCl_2_, 2 CaCl_2_, 5 EGTA, 5 HEPES, and 3 ATP-Na (adjusted to pH 7.2 with KOH). Seal resistances were in the range of 2–5 GΩ.

Superfusion of pre-warmed (36°C) experimental solutions was achieved by using a multi-MPRE8 heating pen (Cell Micro Controls) and data acquisition was performed using Multiclamp 700B patch-clamp amplifiers connected to a Digidata 1322A interface (Molecular Devices, Saint Jose, CA, USA).

### Immunocytochemistry

The DA phenotype of neurons in the process of maturation was confirmed by immunocytochemistry as described earlier ^53^ with the use of tyrosine hydroxylase (TH) (Merck, AB152) and βIII tubulin (TUBB3)/ TUJ1 (BioLegend, MMS-435P) antibodies. Samples were imaged using an inverted confocal microscope (Leica SP5), with a resolution of 512 x 512 pixels, pixel size is 0.758 µm. A ratio of TH+/TUBB3+ was measured by counting the number of TH+ or TUBB3+ cells, in order to assess the maturation of the human mDAN culture ^53^.

### Data analysis

Dynamic-clamp data was digitally filtered at 1 kHz and further exported as MAT-files using Fitmaster (Heka Elektronik). Offline data analysis was performed using custom-written scripts in MATLAB. Spike thresholds were determined using the first derivative of the recording where dVm/dt ≥ 10 mV/ms. Spike thresholds, peaks and minima after hyperpolarization potential were visually verified for each cell. The *g_pace_*-modulated firing frequency was averaged for the first three ISIs (frequency in Hz).

All statistical analysis were performed using GraphPad Prism 10.2.1. Non-parametric Wilcoxon’s tests and Friedman’s test followed by Dunn’s multiple comparison were performed and data are reported as mean ± standard error of the mean (SEM) and as median (interquartile range). Number of cells are denoted by n, number of animals used as N.

## Data and materials availability

All data are available in the main and supplementary text. Codes for simulation can be found at https://github.com/arthur-fyon/Pacemaking_DA_Fyon_2026

## Acknowledgments

We thank our colleagues from the laboratories of Neuroengineering and Neurophysiology for their valuable input and help, especially L. Vandries for the histology of human mDANs. We are grateful to Pr. J-F Liégeois and Dr. R. Vitello for providing the mesylate salt of XG; Pr. C. Canavier & Dr. C. Knowlton for sharing the original code for the pacemaker model; B. Fischer, J. Sonntag, S. Betz, G. Amrhein and T. Wulf for technical assistance in this project; Dr. F.C. Wang, Dr. L. Nguyen & Pr. A. Monteil for their input on the manuscript.

## Author contributions

Conceptualization: AFy, VS, GD & KJ;

Methodology: AFy, PM, MG, SR, JB, AFr, KJ;

Investigation: AFy, OP, NS, PM, AL, MG, & KJ;

Visualization: AFy & KJ;

Funding acquisition: VS, GD & KJ;

Project administration: VS, GD & KJ;

Supervision: JR, MM, VS, GD & KJ;

Writing – original draft: AFy & KJ;

Writing – review & editing: all authors

## Additional information

Supplementary Information is available for this paper

