## Supplementary text and figures for "A new perspective on slow pacemaking in brain and heart"

Supplementary file

### Supplementary Text

*Specific parameter values used*

In **Fig. 5A-B**, the non-specific leak conductance was reduced to 0.01 to lower the firing frequency.

In **Fig. S4**, ten sets of conductances were randomly chosen from the degenerate population with a fixed random seed for reproducibility.

In **Fig. S6**, a NALCN conductance of 0.012 mS/cm$^2$ was used, and values of 0.004 mS/cm$^2$ and 0.012 mS/cm$^2$ were used for the FI curves in **Fig. S6C**. The CaL conductance was reduced to 0.05 mS/cm$^2$ and the Kd conductance to 3 mS/cm$^2$ to enlarge the firing region with respect to the NALCN conductance and facilitate visualization.

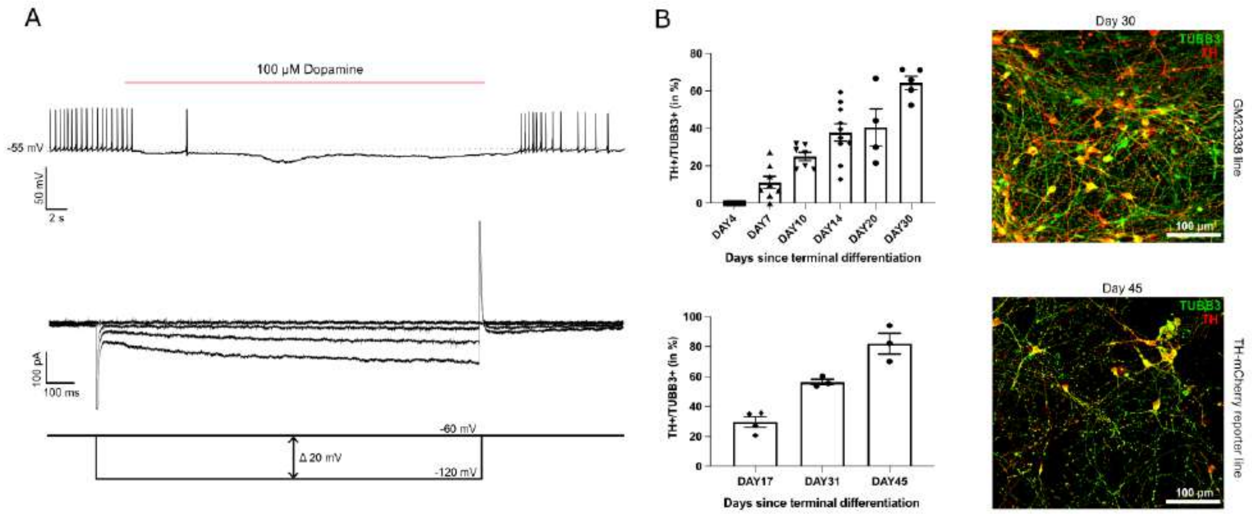


**Fig. S1. Identification of human mDANs**. (**A**) Top panel shows that application of dopamine on mDANs triggers silencing of pacemaking by hyperpolarization due to activation of DA autoreceptors[53], as observed in rodent mDANs[49]. Bottom panel shows an example traces of $I_H$ in human mDANs. (**B**) Evolution of the expression of TH, showing differentiation of neurons into mature mDANs.

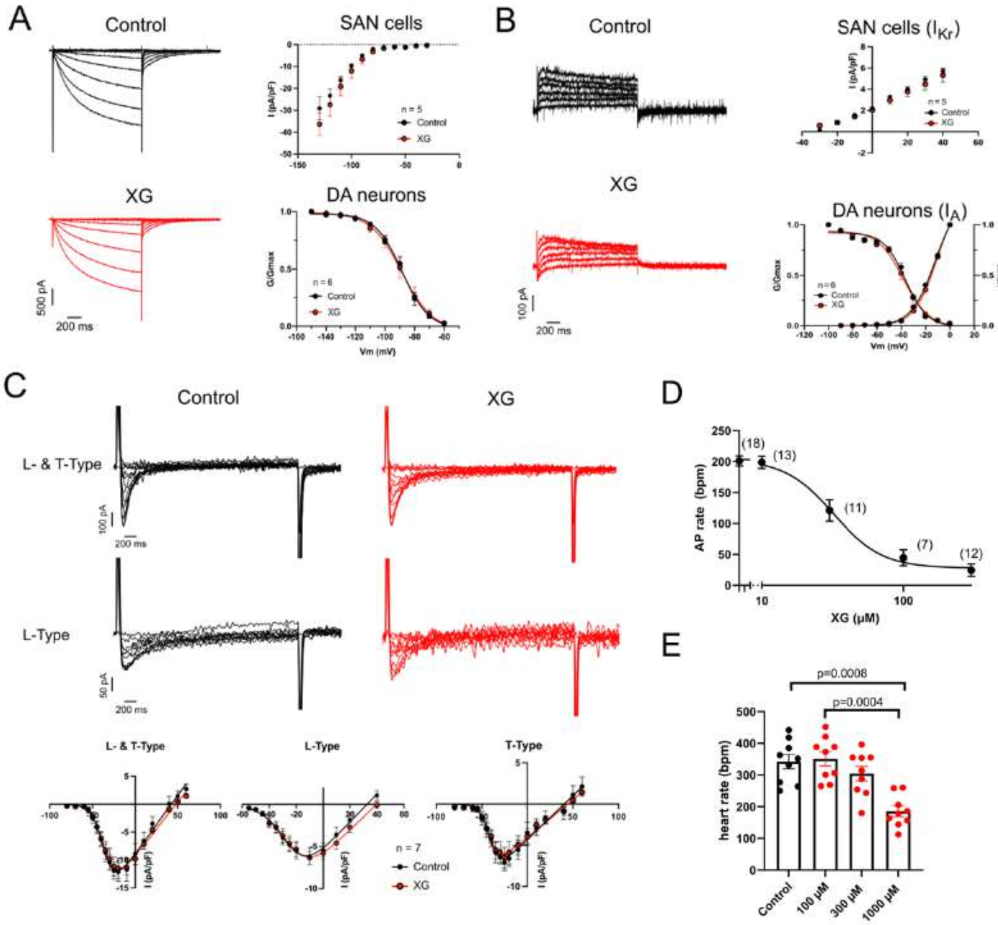


**Fig. S2. Slow pacemaking silenced by XG is not due to an off-target effect on the main voltage-gated channels of SAN myocytes**. (**A**) XG does not interact with neuronal and cardiac HCN currents. The left panels show families of $I_F$ traces activated upon hyperpolarization in SAN myocytes in the absence (black lines), or presence (red lines) of XG (300 µM). The right upper panel show corresponding isochronal current-to-voltage relationships of $I_F$ in the presence (red circles), or absence (black circles) of XG. (**B**) XG does not affect the SAN delayed rectifier ($I_{Kr}$) or A-type potassium channels in DA neurons. The left panels show families of $I_{Kr}$ in SAN myocytes in the absence (black lines), or presence (red lines) of XG (300 µM). The right upper panel show corresponding current-to-voltage relationships of $I_{Kr}$ in the presence (red circles), or absence (black circles) of XG. (**C**) XG does not affect L- and T-type calcium currents in SAN myocytes. The upper panels show families total $Ca^{2+}$ currents ($I_{CaL,}$ L-type and $I_{CaT}$, T-type) and $I_{CaL}$ in the presence (red lines), or absence (Control, black lines) of XG (300 µM). The bottom panels show corresponding current-to-voltage relationships of $I_{CaL}$, $I_{CaT}$ or total $I_{Ca}$ in the absence (black circles), or presence (red circles) of XG. In (**A**) and (**B**), graphs of mDANs were adapted from *Jehasse et al. 2021*[7]. (**D**). Dose-dependence of inhibition of pacemaking in SAN myoctyes by XG. Fitting of dose-response curve yielded half-inhibition ($EC_{50}$) of 32 μM and Hill's coefficient of 2.5. Numbers in parentheses indicate the number of SAN myocytes tested at a given concentration (**E**). Slowing of spontaneous rate in n=9 isolated Langendorff-perfused hearts by increasing concentrations of XG (red circles). Statistics: Friedman's test followed by Dunn's multiple comparison.

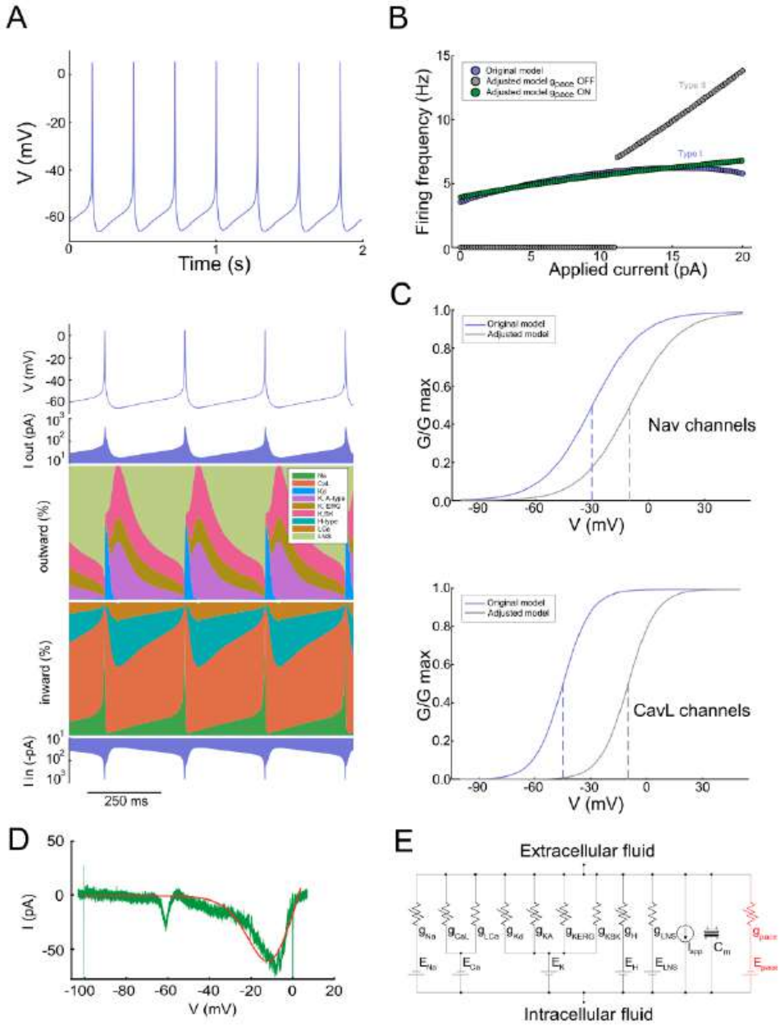


**Fig. S3. Adjustment of current conductance-based model of a mDAN.** (**A**) Simulated pacemaking and currentscape analysis from the model of *Yu et Canavier 2015* [17]. (**B**) Blue dots represent the firing frequency relative to current injections from the model of Yu et Canavier, showing a type I excitability similar to in vitro mDANs. Grey dots represent the firing frequency relative to current injections from the adjusted model of *Yu et Canavier*, changing the excitability type from I to II. Adding $g_{pace}$ to the adjusted model (green dots) shifts the excitability of the neuron from type II to type I. (**C**) Activation curves from $Na_v$ (top) and L-type $Ca_v$ (bottom) channels. Blue curves correspond to the parameters used in the model of *Yu et Canavier*, while grey curves corresponds to our adjustment to physiological values as observed in mDANs for $Na_v$[18] and L-type $Ca_v$ channels[19]. (**D**) Fitting curve (red) based on XG-sensitive current (green) to generate

$g_{pace}$. (**E**) Circuit for conductance-based model with nonlinear conductances and the pacemaking conductance (in red).

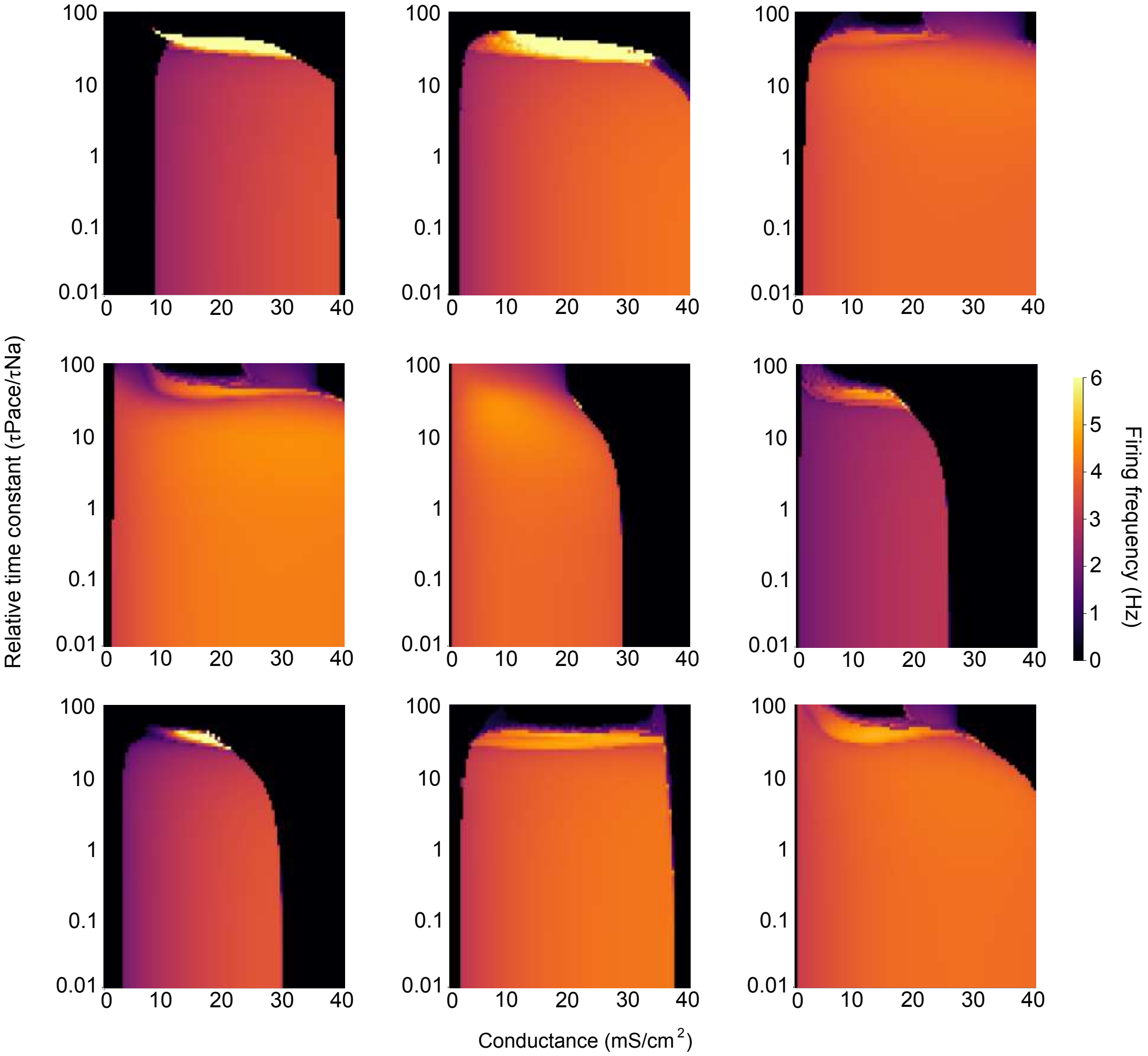


**Fig. S4. Additional heatmaps generated from other random conductance values for all channels.** Overall, we observe a wide range of densities from $g_{pace}$ in order to produce slow pacemaking, with fast kinetics.

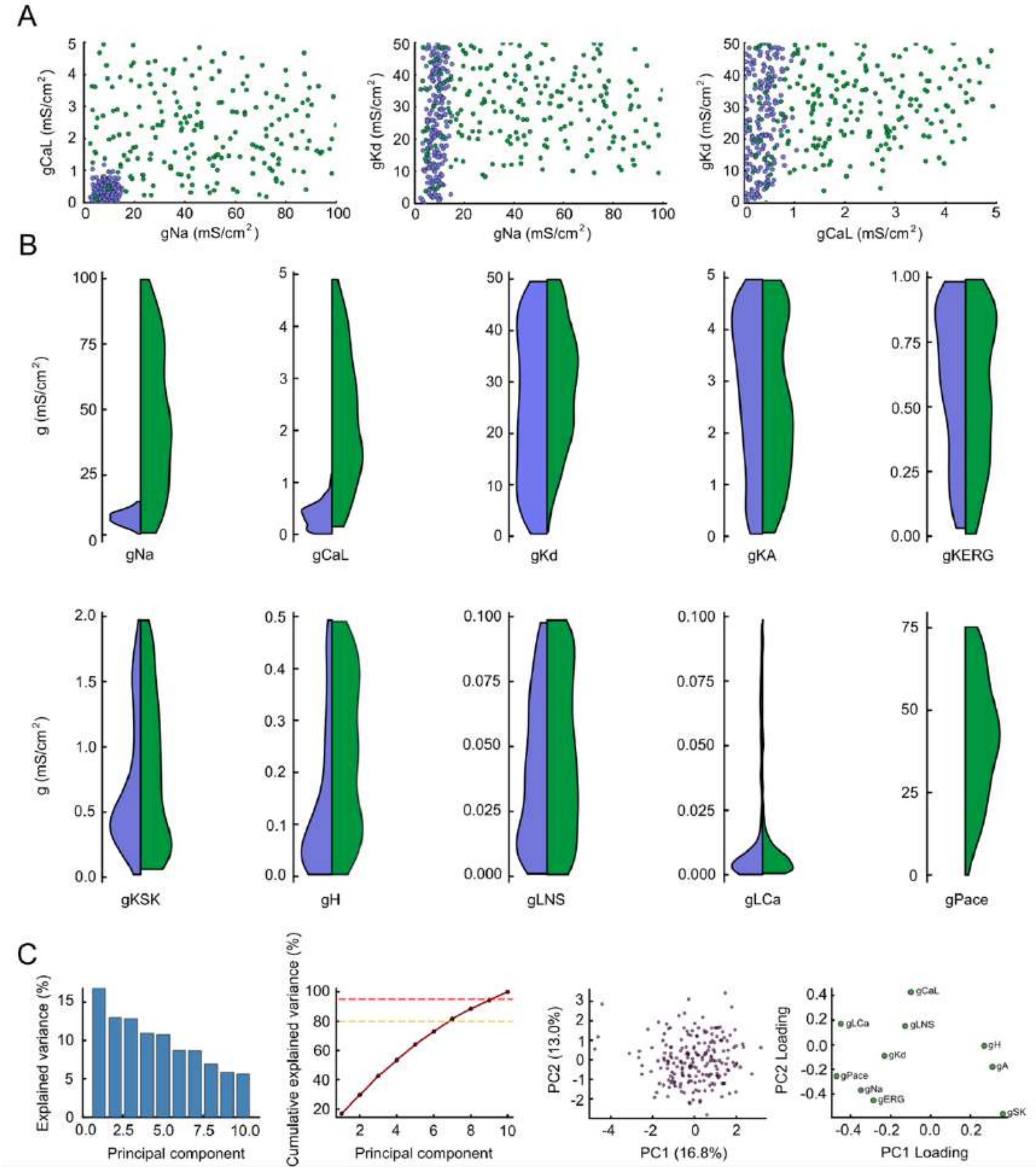


**Fig. S5. Degenerate populations and PCA analysis of the adjusted model.** (**A**) Scatter plots of degenerate populations for the original (blue) and adjusted (green) models in selected conductance subspaces ($g_{Na}$, $g_{CaL}$ and $g_{Kd}$), highlighting that our model allows more degeneracy. (**B**) Violin plots of all conductances for both populations. (**C**) Summary of PCA analysis on the adjusted degenerate population (green in **A** and **B**). From left to right: bar plot of explained variance for each component, scree plot, projection onto the PC1–PC2 plane, and scatter plot of PC1/PC2 loadings.

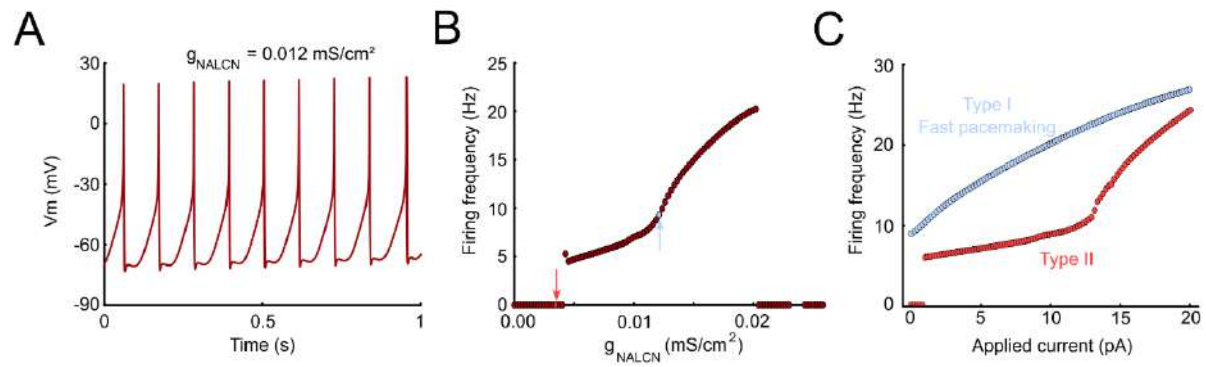


**Fig. S6. Voltage-dependence of $g_{pace}$ is required to generate steady and slow pacemaking.** (**A**) Replacement of $g_{pace}$ by a linear depolarizing conductance in our adjusted model such as $g_{NALCN}$ does not result in slow pacemaking. (**B**) Firing frequency related to the density of $g_{NALCN}$ shows the restricted range of $g_{NALCN}$ density where pacemaking occurs. (**C**) Firing frequency – current relationship curve shows that low density of $g_{NALCN}$ leads to type II excitability, which does not correspond to mDANs excitability, whereas higher density of $g_{NALCN}$ leads to type I fast pacemaking, also not corresponding to mDANs slow firing frequency.
